# DeltaBreed: A BrAPI-centric breeding data information system

**DOI:** 10.1101/2025.04.23.650200

**Authors:** Shawn C. Yarnes, Nick Palladino, David J. Meidlinger, David R. Philips, Heather M. Sweeney, Matthew L. Mandych, Shahana A. Mustafa, Samuel Bouabane, Timothy E. Parsons, Tyler J. Slonecki, Dongyan Zhao, Trevor W. Rife, Bryan J. Ellerbrock, Chaney Courtney, Peter Selby, Mirella Flores-Gonzalez, Lukas A. Mueller, Johan S. Aparicio, Khaled Al-Sham’aa, Sebastian Raubach, Craig T. Beil, Moira J. Sheehan

**Affiliations:** Breeding Insight, Cornell University, Ithaca, NY, United States of America; Plant and Environmental Sciences Department, Clemson University, Florence, SC, United States of America; School of Integrative Plant Science, Cornell University, Ithaca, New York, United States of America; Boyce Thompson Institute, Cornell University, Ithaca, NY, United States of America; Department of Plant and Agroecosystem Sciences, University of Wisconsin–Madison, WI, United States of America; Genetic Innovation Program, ICARDA, Cairo, Egypt; Information & Computational Sciences Department, The James Hutton Institute, Dundee, Scotland, UK

**Author notes:** Corresponding author: (MJS). Current Address: Twilio, San Francisco, CA, United States of America. The Breeding Insight software development team contributed equally to the codebase of DeltaBreed (Shawn C. Yarnes, Nick Palladino, David J. Meidlinger, David R. Philips, Heather M. Sweeney, Matthew L. Mandych, Shahana A. Mustafa, Samuel Bouabane). These authors contributed equally to facilitating interoperability test cases and desired-behavior review (Timothy E. Parsons, Tyler J. Slonecki, Dongyan Zhao, Trevor W. Rife, Bryan J. Ellerbrock, Chaney Courtney, Peter Selby, Mirella Flores-Gonzalez, Lukas A. Mueller, Johan S. Aparicio, Khaled Al-Sham’aa, Sebastian Raubach, Craig T. Beil, Moira J. Sheehan).

## Abstract

DeltaBreed is a unified breeding data management system designed by Breeding Insight (BI, Cornell University) to serve the wide diversity of USDA-ARS specialty crop and animal breeding programs. DeltaBreed has a RESTful microservice architecture that utilizes the BrAPI Java Test Server as its primary database. The system is interoperable with many BrAPI-compliant applications (BrApps), including Field Book, and is continually aligned with the most recent BrAPI specifications. Here, we describe the features of DeltaBreed and provide several interoperability test cases to illustrate successes and challenges encountered during development. We also discuss future expansion and enhancement plans for DeltaBreed, as well as outline possible solutions to known limitations.

## Introduction

<u>DeltaBreed</u> (RRID:SCR_026678) is a unified information system for managing findable, accessible, interoperable, and reusable (FAIR) [1,2] breeding data. Breeding Insight (BI; RRID:SCR_026645) developed DeltaBreed to support the wide diversity of US Department of Agriculture – Agricultural Research Service (USDA-ARS) specialty crop and animal breeding programs. To date, few USDA-ARS specialty breeding programs use a system other than Excel to manage different data types, such as germplasm, pedigrees, observation variables, experiments, observations, genotype samples, and genotyping results. USDA-ARS specialty breeding programs encompass considerable logistical efforts managed by small research teams responsible for trialing, advancing, and selecting improved germplasm. Funding, staffing, and time constraints [3] place these programs at a data management disadvantage. Breeding data are characterized by disparate data types and increasingly large data volumes that are difficult to store, query, join, and analyze without support and infrastructure for server-based solutions. BI strives to fill this gap by developing and administering the DeltaBreed web application for USDA-ARS partners.

DeltaBreed v1.0 is a minimum viable product: the most streamlined species-agnostic system required to manage breeding data originating from BI’s <u>collaborating breeding programs and germplasm repositories</u> [4]. Breeders have shared substantial phenotypic and pedigree data with BI to assist in the validation of genotyping panels [5–8], support marker-trait associations, and perform population structure analysis [9–11]. We and others [12] have found that data from phenotyping experiments are often ambiguous due to undefined variables, murky pedigrees, and missing identifiers. DeltaBreed v1.0 was built as a foundation to ensure that breeders moving forward can intuitively maintain FAIR breeding data that supports compatibility and interoperability with apps developed by BI and others, as well as the public.

DeltaBreed is an open-source application (Apache License, v2) that uses the Breeding Application Programming Interface (BrAPI), a community-defined standard [13], as the primary way to transmit breeding data between different software applications. The BrAPI technical specification provides standardized API definitions for DeltaBreed to successfully communicate data between a suite of BrApps (BrAPI-compliant Applications). The release of DeltaBreed demonstrates a BrAPI-centric approach that leverages open-source software from unrelated or loosely related initiatives. While not yet fully realized, seamless interoperability between BrApps is now an attainable goal.

## Results

### System architecture

DeltaBreed has a RESTful (Representational State Transfer) microservice architecture (Fig 1), which differentiates it from other open-source breeding information systems that employ single-tier application frameworks [14–17]. Instead of building the system around a single database, BI has taken a modular approach, using REST APIs (primarily BrAPI), to communicate data between web services. The advantage of modularity is that services can easily be added, updated, or swapped as needed and with minimal impact on users. Optimizing performance is one challenge of the RESTful microservice approach. The latency between systems and the architecture of individual services impact overall performance, which can be exacerbated by poor internet connectivity. DeltaBreed v1.0 employs a Redis cache to reduce user interface (UI) load times.

**Fig 1.**
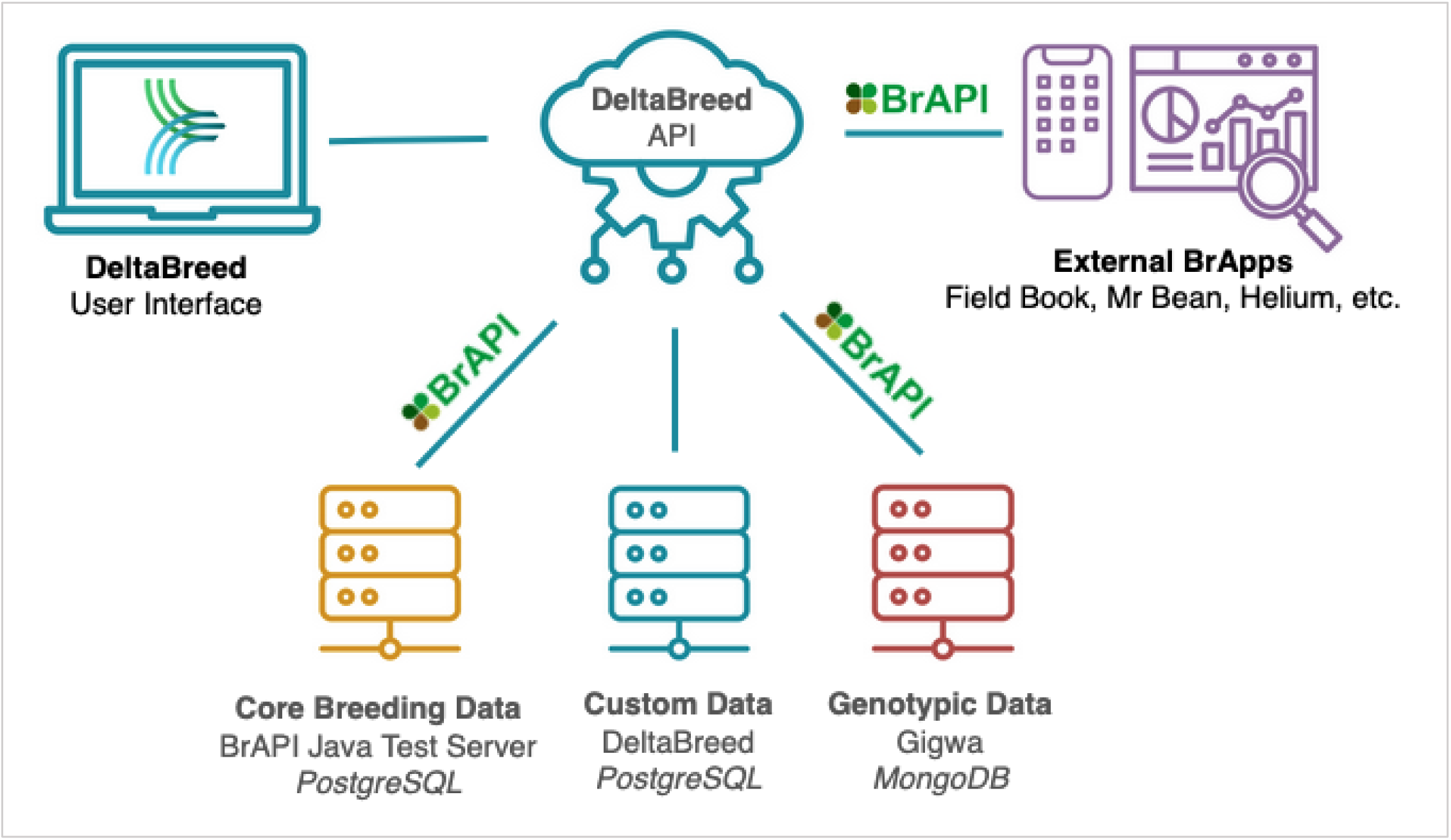
DeltaBreed’s Modular Architecture Users can interact with DeltaBreed data through the web interface and via external BrApps. REST APIs, primarily BrAPI, are used to communicate breeding data among various web services, including DeltaBreed databases. DeltaBreed communicates with two BrAPI databases and uses a custom database to handle data, not supported by BrAPI, like user management. Disparate datatypes are handled by connected subsystems with optimized data architectures suitable to the data storage needs.

#### Interoperability test case 1: BrAPI Java Test Server & BreedBase

The DeltaBreed system requires a fully BrAPI-compatible database for storing breeding data, specifically BrAPI-core, BrAPI-germplasm, and BrAPI-phenotyping data. Breeding Insight has successfully integrated with two such databases, BreedBase [15,18] and the <u>BrAPI Java Test Server (BJTS).</u> BreedBase is a standalone system with a single-tier architecture (2.3M lines of code) that uses BrAPI mainly for communication with external apps. The BJTS is a fully implemented example of the BrAPI specification, designed for the BrAPI community to test the implementation of BrAPI endpoints [19]. As the BJTS was previously untested, BI initially made use of BreedBase to store DeltaBreed production data. While DeltaBreed and BreedBase are interoperable via BrAPI, we achieve shorter load times when DeltaBreed is connected to the lighter-weight BJTS (0.5M lines of code). DeltaBreed v1.0 now utilizes the BJTS as its primary database. This test demonstrates that a microservice architecture supported by BrAPI permits changes to the database schema that would be more difficult in a single-tier application.

#### User Experience (UX)

Despite having modular architecture, DeltaBreed provides a unified UX for managing diverse data types. Users can learn DeltaBreed features without being aware of BrAPI or the underlying services. The initial users of DeltaBreed v1.0 are BI coordinators and data curators who support USDA-ARS partner programs adopting FAIR data management practices. BI coordinators have been instrumental in ensuring that DeltaBreed is easy to use and well-documented by providing comprehensive UX testing, a <u>user manual,</u> and <u>training materials</u>.

DeltaBreed establishes data standards and validations previously absent in USDA-ARS partner programs. Common to all DeltaBreed data management modules are simple workflows, defined data requirements, informative error messaging (Fig 2), transparent data views (Fig 5), and cohesive formatting of template and download files.

**Fig 2.**
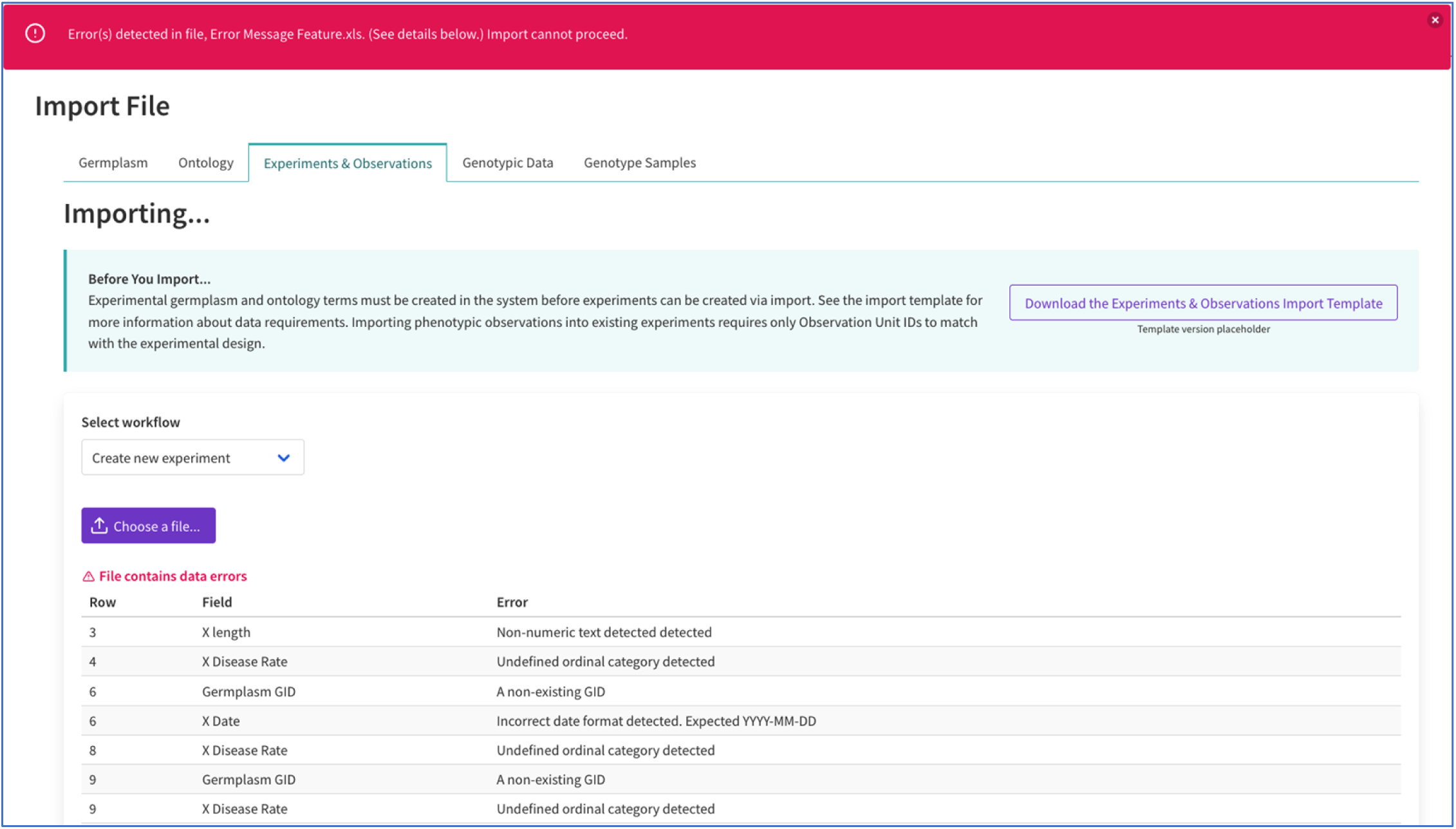
DeltaBreed file import UI This figure illustrates DeltaBreed’s clear and actionable UI error messaging. In this example, there are several data errors detected in the Experiment and Observation import file. An error table alerts users to the exact row in the import file that needs correction.

**Fig 3.**
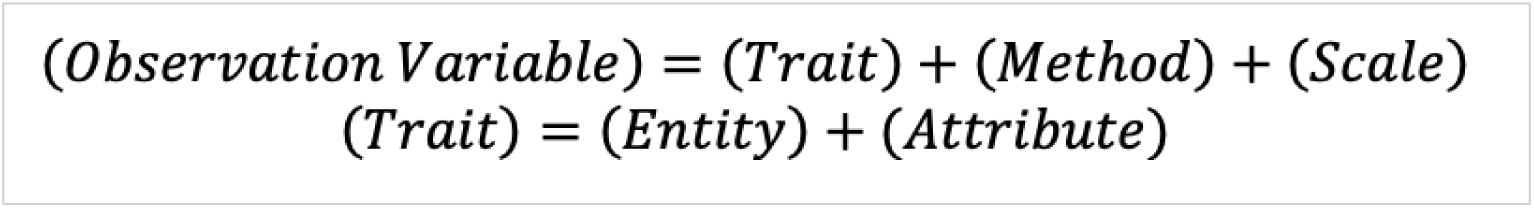
Observation variable conceptual module adapted from Crop Ontology

#### Authorization and program privacy

DeltaBreed helps the USDA meet its open data requirements by balancing privacy, security, and openness. DeltaBreed is a private system administered by BI. Invited users authenticate themselves using their <u>ORCID</u> <u>(Open Researcher and Contributor ID)</u>, a digital identifier that integrates with DeltaBreed through the open authorization (<u>OAuth 2.0</u>) protocol. Users interact with breeding data from within their associated program(s). Different program roles and permissions allow for various levels of data openness within a program, including program administration, read-only, and experimental collaboration. Data openness is entirely controlled by breeding program administrators and can be employed at various levels to suit the lead breeders’ needs.

### Ontology

All breeding information systems have some form of controlled vocabulary (i.e., ontology). The DeltaBreed Ontology module allows users to create custom observation variables and establish standards for observation validations. DeltaBreed defines observation variables following the <u>Crop Ontology</u> conceptual module [20,21] standards and validations for observations. For example, DeltaBreed observations defined by numerical variables are validated not to contain alphabetic characters, and observations for text variables are not validated at all (Table 1, Fig 2).

**Table 1.** Observation variable scale class and corresponding observation validation.

| Observation variable scale class | Observation data validation |
| --- | --- |
| Numerical | Numbers only; no alpha characters |
| Date | Date in YYYYMMDD format |
| Ordinal | All categories or values are defined |
| Nominal | All categories are defined |
| Text | none |

### Germplasm Management

The DeltaBreed concept of germplasm, by design, is very broad. Germplasm is defined as any biological unit (gamete pool, individual, clone, or population) capable of reproduction via any breeding method (DeltaBreed offers >60 pre-defined methods with the option to add customized ones). Drawing from the ICIS Genealogy Management System model [22], each DeltaBreed germplasm record is linked by breeding method to its progenitors through unique IDs. DeltaBreed assigns both UUIDs (Universally Unique IDs) and human-readable sequential numbers (germplasm IDs, GIDs) to every germplasm record. Dependence on unique IDs alleviates the need to adhere to complex naming logic to convey germplasm details, including pedigree connections, and allows for duplicate named entities that are, in fact, separate records (a common occurrence in clonally propagated species). Names make poor unique identifiers because they often break down due to common spelling mistakes, symbol use, and the tendency of names to evolve over time, such as during the selection process and at variety release. Names are also unsuitable for uniquely differentiating clonal germplasm, which commonly inherits the name of its progenitor. For example, grape varietal clones are generally uniquely named only after the discovery of phenotypically important somaclonal variation. To support breeding for any species, DeltaBreed accepts all germplasm naming conventions, permits name duplicates, and allows for custom breeding methods.

#### Interoperability test case 2: UI integration of BrAPI pedigree viewer

Genealogical records are universal to all breeding programs, and breeders want to visualize relatedness in pedigree trees. We have fully integrated an interactive pedigree viewer into the DeltaBreed by adapting a previously-existing open-source BrAPI Pedigree Viewer developed by the Boyce Thompson Institute [24]. In this case, we found that integrating this BrApp was faster than developing a comparable feature *de novo*. Most BI development focused on refining and enhancing the visualization in the DeltaBreed UI. We added the GID to the germplasm name and developed an unknown germplasm placeholder, “Unknown [GID: 0]” (Fig 4A). Pedigree unknowns, especially for the male parent, are relatively common in pedigree records. DeltaBreed handles “Unknown [GID: 0]” differently than other germplasm records, in that it can appear multiple times in a pedigree view, avoiding the misleading impression that a single record is making an outsized genetic contribution to the program or tangling pedigree connections (Fig 4A).

**Fig 4.**
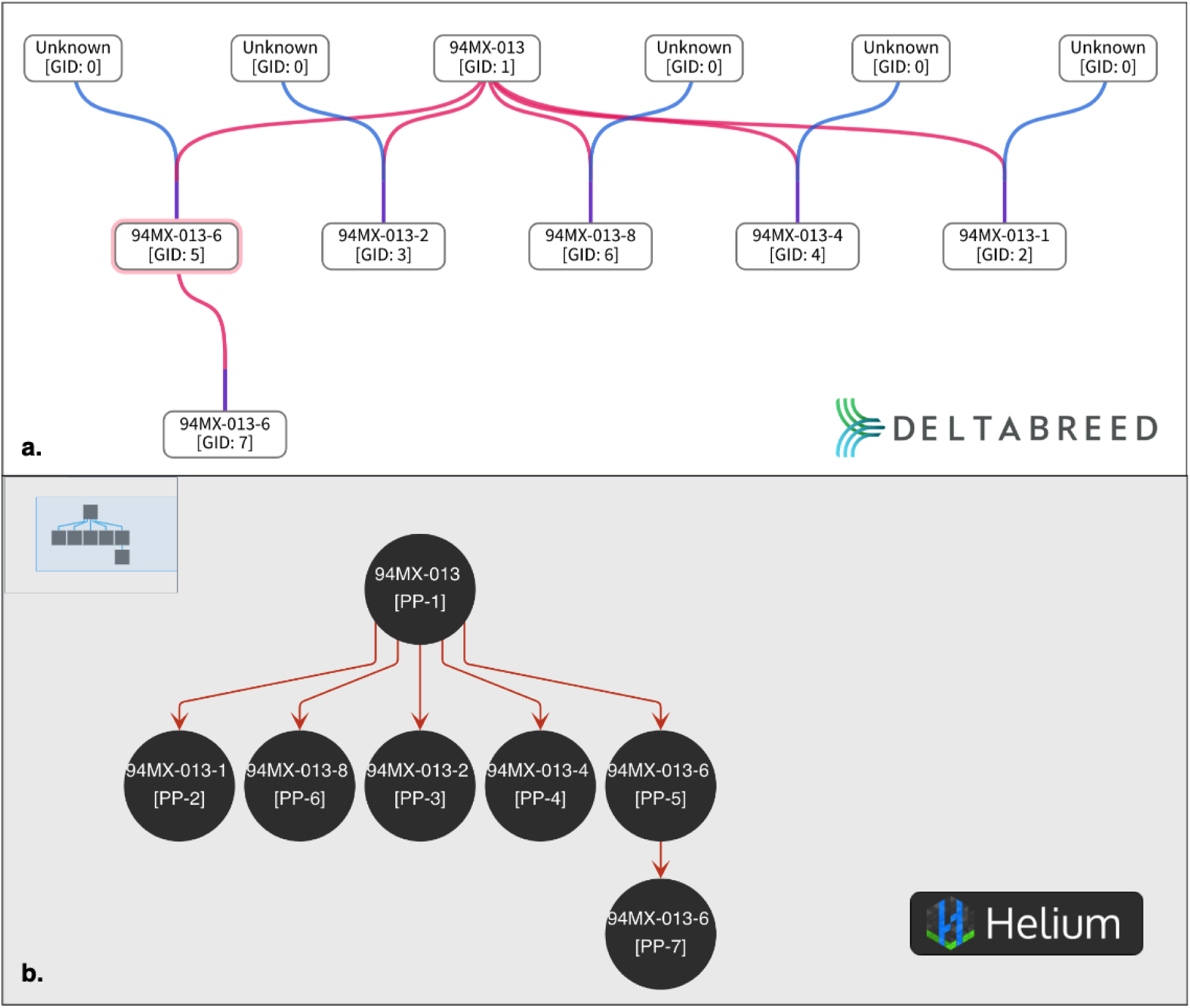
Pecan Pedigree View Comparison This figure represents the fully expanded pedigree of 94MX-013-6 [GID: 5] using the integrated DeltaBreed viewer (A) and Helium (B). (Genealogy Courtesy of Warren Chatwin, USDA-ARS Pecan Somerville, TX)

#### Interoperability test case 3: Helium pedigree viewer BrAPI connection

<u>Helium</u> is a pedigree viewing BrApp developed by The James Hutton Institute (JHI) [25,26] that includes commercial libraries incompatible with DeltaBreed’s Apache 2 license. Although license incompatibility limits our ability to integrate Helium fully, we recently collaborated with JHI to allow Helium users to authorize and pull pedigree data directly from a DeltaBreed program. DeltaBreed users can use Helium filter capabilities, specifically filter by accession number (DeltaBreed GID), to visualize the genealogy of a single germplasm record (Fig 4B).

However, limits to interoperability remain. DeltaBreed programs with more than a few thousand germplasm records cannot use Helium’s default pedigree retrieval because thousands of pedigree nodes are not meaningfully visualized in a single view, and load times are prohibitive for practical use. Additionally, Helium and other BrApps. DeltaBreed displays a simple germplasm name in the UI but saves germplasm names to the database as a concatenation (i.e., name + program code + GID) to ensure compatibility with BrApps that do not differentiate data by programs and require unique germplasm names. Although derived from DeltaBreed, the program code and GIDs rendered in the Helium UI (see rigid brackets in Fig 4B) are not details a DeltaBreed user would expect or desire to be included in a germplasm name and appear as an artifact of interoperability. Also, note that Helium does not recognize DeltaBreed’s “Unknown [GID: 0]”, in part because there is not a BrAPI mechanism to transmit the special case of an “unknown” germplasm record.

### Experiments and observations

DeltaBreed v1.0 allows users to create and append single or multi-environment experiments. Experiments describe the spatiotemporal arrangement of germplasm (GIDs) into observation units (ObsUnitIDs) to make observations and detect statistical differences in phenotypic response. Experiments can also describe non-randomized arrangements, such as nurseries or crossing blocks, that may have purposes beyond phenotyping. The DeltaBreed experiment concept aligns closely with Minimal Information About a Plant Phenotyping Experiments (<u>MIAPPE</u>) standards [27], including most terminology and BrAPI mappings.

Active experimental management in DeltaBreed is a multi-step process that begins with creating an experiment prior to data collection (Fig 5) and ends with appending observational data. Interoperability test cases 4 and 5 both involve active experimental management using the <u>Field Book</u> mobile app [28,29]. Field Book is a widely used BrApp for recording phenotypic observations and measurements on handheld Android devices like phones and tablets. Field Book is part of the PhenoApps project, and BI has worked closely with the PhenoApps team to improve its BrAPI interoperability.

**Fig 5.**
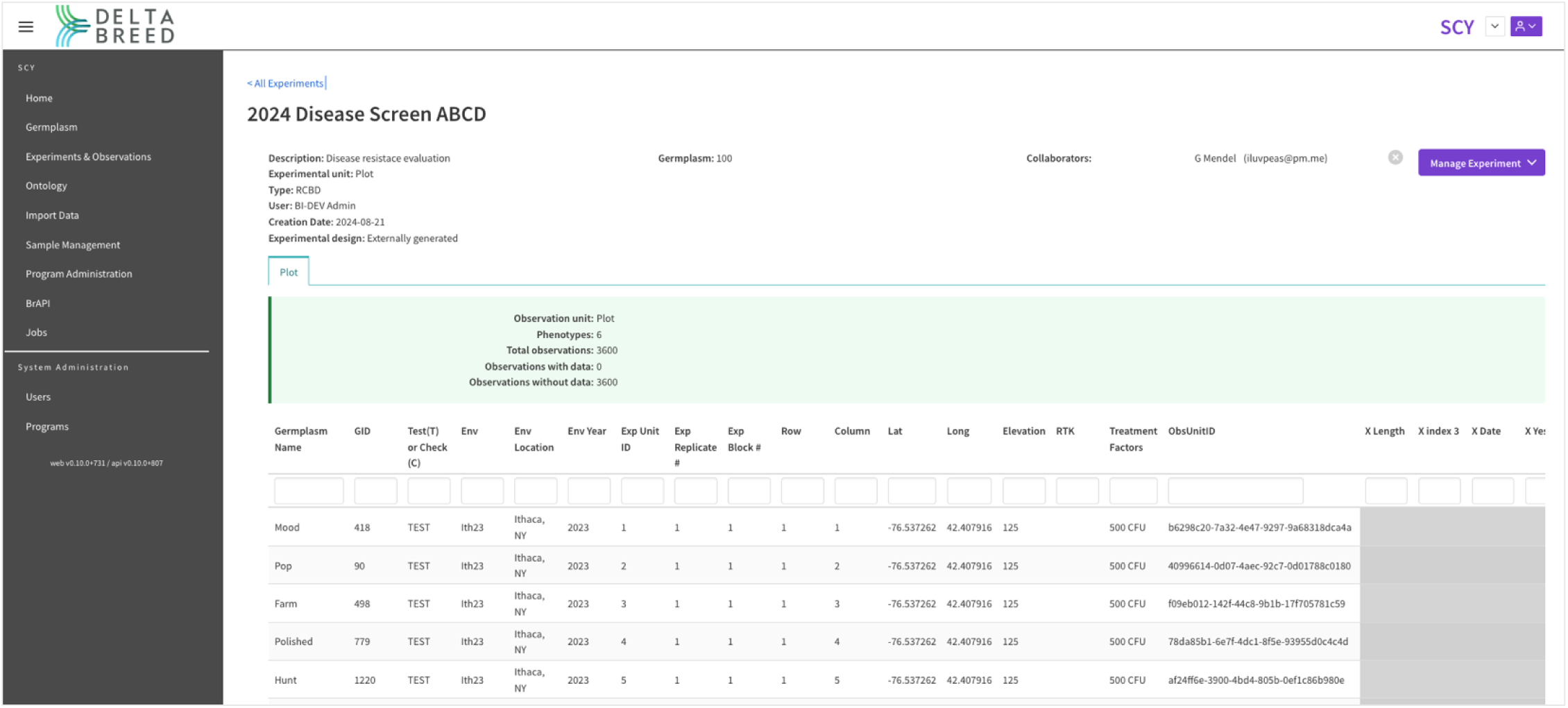
DeltaBreed experiments and observations user interface This screenshot displays an experiment before data collection. Each table row represents an observation unit with its unique ObsUnitID and associated human-readable germplasm record (GID). The grey cells represent pending observations. Experimental observation variables (e.g., “XLength”) are defined in the DeltaBreed Ontology module.

#### Interoperability test case 4: DeltaBreed, BreedBase, and Field Book

Before the completion of the phenotyping workflow in DeltaBreed, BI integrated several BrApps to support active 2021 experimental data collection by USDA-ARS blueberry breeders. We used the DeltaBreed UI to establish observation variables, which were stored in BreedBase, and utilized the BreedBase UI to establish studies before data collection. We also enabled Field Book to authenticate and pull studies and observation variables from BreedBase and push observations made in Field Book back to BreedBase after data collection. This effort served as a successful proof of concept for multi-application BrAPI integration but highlighted limitations in the process of accepting observations back without quality control validations. Manual curation was ultimately required to resolve missing, duplicated, and erroneous observation values arising from inadvertent partial observation uploads, repeated duplicated uploads by single users, and extensive uploads to the database by multiple data collectors on multiple devices.

#### Interoperability test case 5: DeltaBreed & Field Book

BI developed new features to support the DeltaBreed-Field Book data collection workflow. Field Book users can authenticate and pull DeltaBreed experimental data collection details to their handheld device without using flat files. The main advantage of this workflow is that observations made in the Field Book app are

DeltaBreed programs (Fig 6A) to support Field Book’s auto-configuration (Fig 6B), eliminating the need to configure BrAPI settings on each tablet by every user manually. The process of selecting observation variables for data collection in Field Book was also streamlined. Observation variables associated with DeltaBreed experimental datasets are now automatically selected in Field Book, reducing the need to select variables in the field manually. DeltaBreed v1.0, like BreedBase (Test Case 4), lacks adequate transaction handling to support BrAPI data pushes from external apps. To avoid time-consuming resolution of data discrepancies encountered in the previous test case, DeltaBreed v1.0 does not accept data from external apps via BrAPI. Observations are uploaded by flat file, ensuring a human-mediated QC process for appending an experiment with observations. The 2024 field season included USDA-ARS alfalfa and hop beta testers, with both groups reporting successful data collection activities (Zhanyou Zang and Tyler Gordon, personal communications).

**Fig 6.**
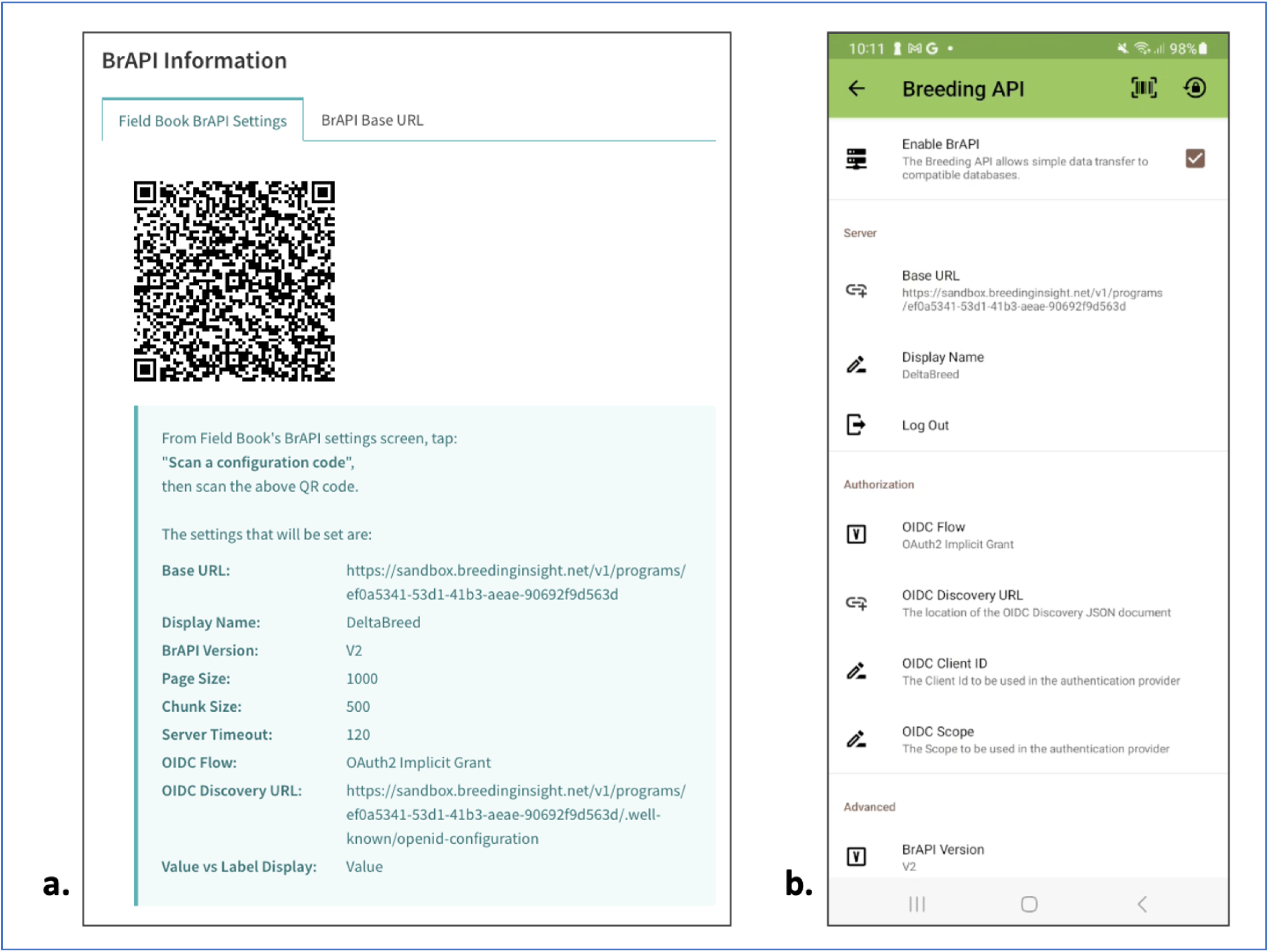
DeltaBreed BrAPI interoperability with Field Book This figure shows side-by-side screenshots of (A) DeltaBreed’s Field Book BrAPI Settings and (B) Field Book’s BrAPI settings after auto-configuration. DeltaBreed auto-configuration is needed to support pulling experimental datasets and observation variables in Field Book for data collection.

#### Interoperability test case 6: MrBean & QBMS

MrBean is an R-Shiny app designed for users with no R programming experience to visualize and model spatial trends in single and multi-environment experiments [30,31]. MrBean works with a core set of R packages, including the QBMS package [32,33], which allows MrBean to access BrAPI endpoints. In collaboration with MrBean and QMBS developers, BI has updated MrBean and QBMS code bases to allow users to authenticate to DeltaBreed and pull experiment and observation data via BrAPI. We have successfully used MrBean to generate descriptive statistics and plots (Fig 7) and perform *lme4* model fitting [34] on DeltaBreed data. Additional effort is needed to integrate other MrBean modeling options.

**Fig 7.**
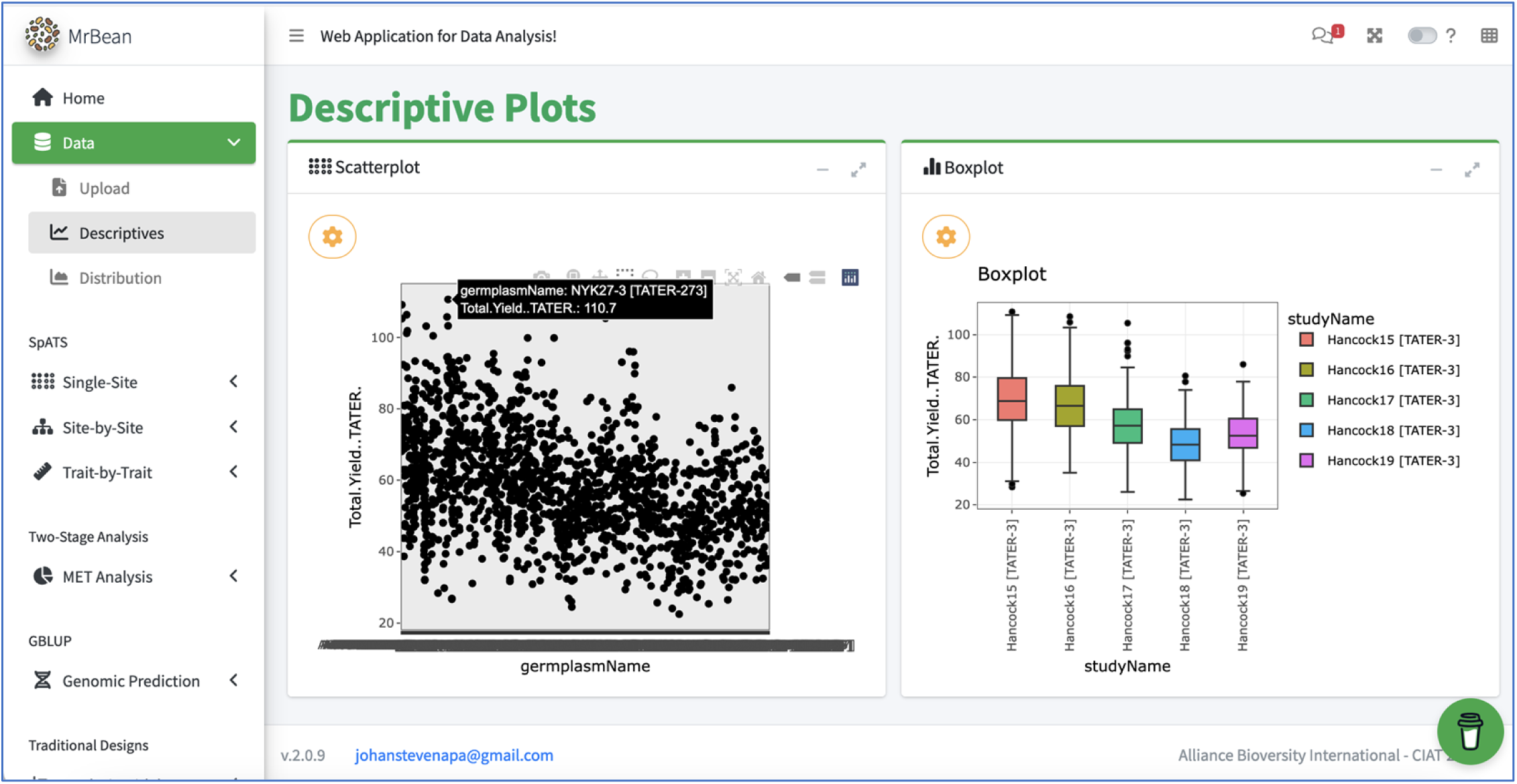
MrBean descriptive plots using potato data pulled from DeltaBreed via BrAPI Notice in the scatter plot that the selected germplasmName is appended with program code & GID, [TATER-273]. Similarly, studyName in the boxplot is appended with program code and experiment ID. These appends are hidden from users on the DeltaBreed UI and are artifacts of DeltaBreed interoperability logic, also described with Helium integration (see Fig 4).

### Genotype Sample Management

DeltaBreed’s sample management module supports the BI genotyping sample submission process. BI has facilitated the genotyping of ∼71K samples for our partner programs through a multistep process that consists of collecting sample submission details, submitting the genotyping order to the vendor, and ultimately receiving and processing the SNP results back from the vendor. Essential to this process are DeltaBreed-generated unique IDs compatible with all systems involved, including human-readable sample names. Samples can be associated with either germplasm records (GIDs) or experimental observation units (ObsUnitIDs), the latter of which is more information-dense. Observation units connect samples to germplasm records, spatiotemporal details, and phenotypic observations.

#### Interoperability test case 7: DeltaBreed and Gigwa

BI is actively developing a complete genotyping workflow and has implemented a DeltaBreed prototype that fully integrates The Genotype Investigator for Genome-Wide Analysis (Gigwa, RRID:SCR_017080; https://southgreen.fr/content/gigwa) into the DeltaBreed architecture (Fig 1) for genotypic data storage [35–37]. The prototype is a proof of concept demonstrating that Gigwa can support DeltaBreed genotypic data loading (VCF data) and microhaplotype sequence visualization (Fig 8).

**Fig 8.**
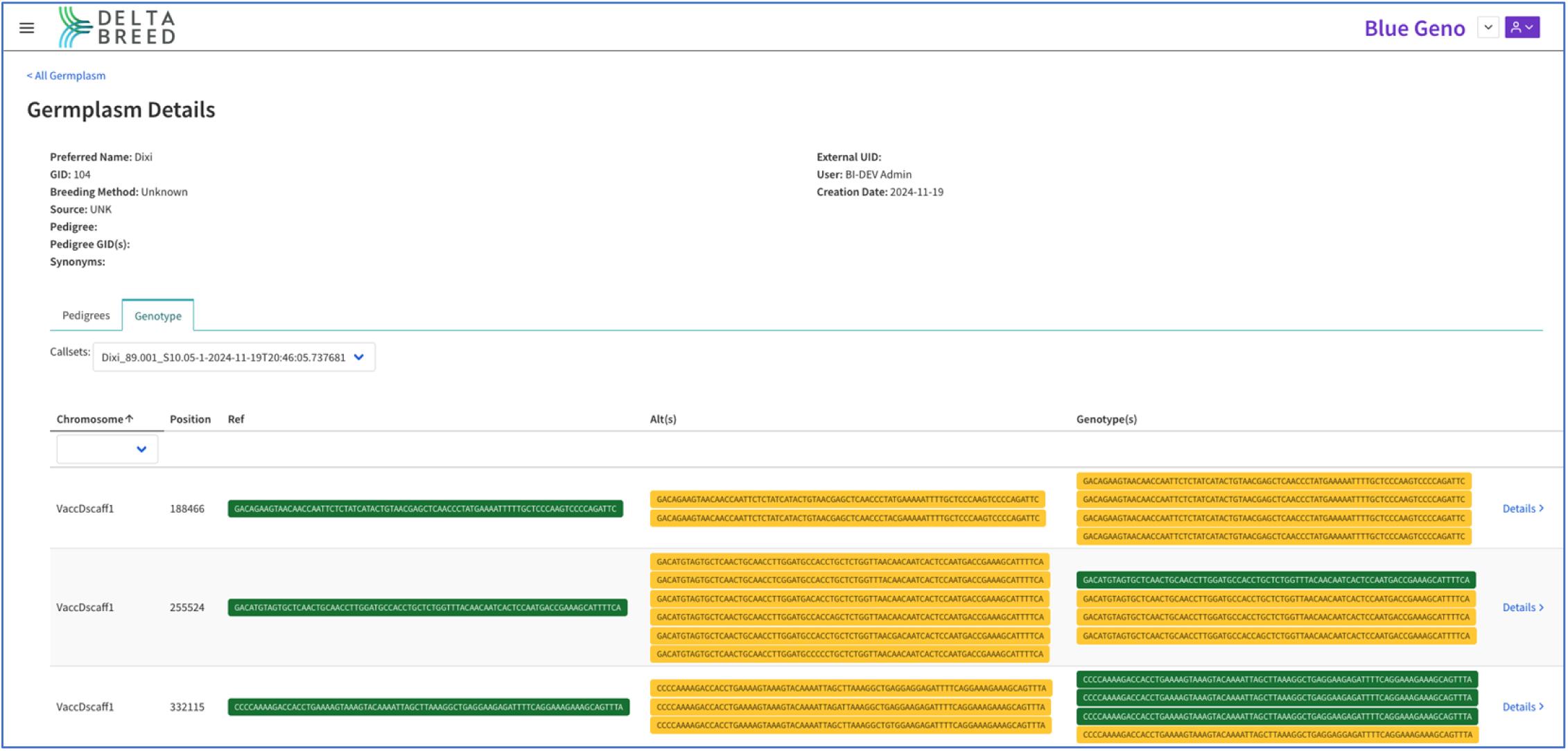
Germplasm microhaplotype view An autotetraploid blueberry (2n=4x=48) germplasm record with BI-processed and validated results for three DArTag loci. At each marker locus, a microhaplotype represents a 54-81bp amplified sequence containing the target SNP variant site (column “Ref”) and/or off-target variants (column “Alt”). In this example, the ‘Dixi’ cultivar harbors four copies of “Alt” alleles at locus VaccDscaff1_188466, one copy of the “Ref” and three copies of the “Alt” alleles at locus VaccDscaff1_255524, and three copies of “Ref” and one copy of the “Alt” alleles at locus VaccDscaff1_332115.

## Discussion

DeltaBreed v1.0 demonstrates the power and unique development challenges of open-source community models and standards in the deployment of FAIR breeding data management tools. DeltaBreed interoperability test cases confirm that a BrAPI-centric microservice architecture is a nimble approach to incorporate services or swap underlying databases (Test Case 1). Open-source BrApps with compatible licenses can be fully integrated into DeltaBreed with less development effort than *de novo* coding (Test Cases 1, 2, 4, 5, 7). While any BrApp is theoretically interoperable with DeltaBreed, we have found that interoperability with independently developed and maintained BrApps is not guaranteed, and even when the connection is technically possible, the user experience benefits from additional refinement (Test Cases 1-6).

BrAPI is integral to DeltaBreed. In addition to facilitating interoperability with independent BrApps, BrAPI facilitates DeltaBreed’s intersystem connections. The BrAPI Java Test Server has proven to be a modestly performant breeding database for DeltaBreed v1.0. The BI and BrAPI teams are actively collaborating on creating a BrAPI Java Production Server (BJPS) with enhanced performance, full CRUD functions (create, read, update, and delete), and transaction handling. Future versions of DeltaBreed will utilize the BJPS and continue to be aligned with the latest version of the BrAPI specifications. Future DeltaBreed features, including new delete functions, will be possible with BrAPI 2.2 updates.

DeltaBreed interoperability is the result of the BrAPI community cross-initiative collaboration. Every DeltaBreed interoperability test case described in this manuscript required BI development effort and most prompted ongoing and continuous collaborative efforts with other software initiatives, ranging from bug reporting, to feature requests, code-sharing, and co-working at annual BrAPI Hackathons. We have found that collaborative solutions tend to be most efficient when collaborators benefit by gaining new features and/or new users while remaining focused primarily on their own code bases and development priorities. Collaborative solutions also tend to be generalized to the benefit of the larger breeding community. For example, a DeltaBreed use case may have prompted the implementation of OAuth protocols in Field Book and Helium, but now that the protocol is in place, it is available to any system.

Systems, like Deltabreed and others, that warehouse breeding data have complex and largely unavoidable differences in business logic. Differences, particularly in naming rules, pose challenges to the interoperability of community BrApps designed for data collection, analytics, and decision-making. For a BrApp to be interoperable with any system, it needs to be flexible and independent of systems logic. We recommend that community BrApps use BrAPI DbIds to detect duplicates if needed, rather than name properties of BrAPI objects. We also recommend that community BrApps provide information-rich UIs, including a complete representation of the available BrAPI object properties, especially when providing search or filtering capabilities. More information improves everyone’s user experience and helps support interoperability. For example, DeltaBreed users need multiple BrAPI properties to uniquely identify germplasm (germplasmName, assessionNumber) and experimental datasets (trialName, studyName, observationLevel).

BI is actively using, maintaining, and improving DeltaBreed in a continuous integration and continuous deployment framework. The system is now available to select USDA-ARS specialty breeding programs. Beta testers have responded positively, especially to interoperability with Field Book. Even when interoperability is not fully UX optimized (Test Cases 4 & 5), our testers enthusiastically adopted Field Book data collection workflows. Continued improvement of DeltaBreed interoperability and UI/UX with Field Book and other BrApps is a priority for BI. We are working closely with the PhenoApps team and others to improve and expand processes. However, breeding terminology and human language remain usability sticking points.

Related BrApps may agree on BrAPI communication standards, but they rarely agree on the terminology in their UIs. For example, an “experimental dataset” in DeltaBreed is equivalent to a “field” in Field Book and a “studyName” in MrBean. As BI looks to the future, we are exploring options to make external BrApps appear more seamless with DeltaBreed without interfering with the UI/UX of non-DeltaBreed workflows. In addition to UI/UX improvements, we are developing support channels and step-by-step, workflow-specific training materials for breeders interested in using the system.

The release of DeltaBreed v1.0 provides USDA-ARS specialty crop and animal breeders with a species-agnostic, open-source data management system with a friendly UI/UX and low barriers to adoption. The web application makes it easy for users of any operating system to migrate data currently stored in Microsoft Excel spreadsheets into a robust repository with data backups, validations, and privacy. Breeders of specialty species, particularly perennial, clonal, or livestock species, will find the flexibility they need in DeltaBreed that is not generally found in software built with only annual row crop breeders in mind.

DeltaBreed has the potential to accelerate delivery of improved genetic cultivars to US farmers and producers by providing the data standardization required to leverage USDA-ARS phenomic and genomic breeding data fully. DeltaBreed is open source and available for any organization to deploy to the infrastructure of their choice. DeltaBreed’s modular nature and BrAPI-centric design should make the system attractive to universities and small to mid-size breeding companies who want to standardize their data without reinventing a core platform, while maximizing the customization possible via BrAPI interoperability. Planned upgrades to the BJPS will deliver an enterprise system suitable for managing large-scale operations, such as the breeding effort across all USDA-ARS crops and animals (>100 crops and ∼15 animal species). USDA-ARS national programs invested 1.0 Billion dollars in improving crops and food animals through Crop Production and Protection (CPP) and Animal Production and Protection (APP) [38]. Also in 2023, the crop returns to U.S. farmers across all commodities combined were more than 109 Billion dollars, a return-on-investment greater than 1:100 per taxpayer dollar [39]. Moreover, most of these species do not have private-sector genetic improvement enterprises, leaving USDA-ARS as a significant single entity for crop and livestock improvement. The consolidation of all breeding management data across the USDA mission areas (which would include the National Arboretum and US Forest Service, both of which engage in breeding efforts) would significantly unify data collection, ensure data integrity and security, and improve cross-collaborations within the agency for a fraction of the cost that other software-as-a-service systems currently available charge. Lastly, DeltaBreed’s modular architecture and BrAPI interoperability provide limitless potential for breeding workflow customization and full integration of diverse data types into routine breeding decisions.

## Author Contributions

SYC documented all user test-case scenarios for DeltaBreed development, approved all acceptance criteria, provided coordination of development efforts, wrote the DeltaBreed manual, and wrote this manuscript. NP, NKJ, DRP, HMS, and MLM, developed and contributed equally to the DeltaBreed code and features. SAM managed and completed all Quality Assurance testing for DeltaBreed. TEP and SB wrote technical specifications for DeltaBreed code, performed code review, and managed the developer team at Breeding Insight. TWR, BJE, and CC provided access and support in the interoperability efforts between Field Book and DeltaBreed via BrAPI. PS is the global BrAPI coordinator and provided support in DeltaBreed’s adherence to the BrAPI specification, provided access and maintenance to the BrAPI Java Test Server, and has organized and hosted numerous Hackathons from which some interoperability solutions included in this manuscript came. MFG provided technical changes to BreedBase to implement or enhance its BrAPI endpoints for improved interoperability. LAM provided BreedBase to the BI team and managed the BrAPI compliance efforts by MFG. JSA is the creator and owner of MrBean and assisted directly in testing the interoperability of DeltaBreed and MrBean. KAS is the creator and owner of QBMS and assisted directly in testing the interoperability of DeltaBreed and QBMS. SR assisted directly in testing the interoperability of DeltaBreed and JHI’s Helium software. CTB provided feature refinement and managed user acceptance testing of all DeltaBreed features. MJS provided funding, project management and supervision, and the writing and revising of this manuscript.

## Acknowledgements

The authors thank Dr. Alexandra Casa for her careful review of this manuscript and for her valuable suggestions for its improvement.

## Funding Sources

### Breeding Insight at Cornell University

These materials are based upon efforts from Breeding Insight (RRID:SCR_026645), a USDA-ARS initiative hosted by Cornell University (supporting SYC, NP, DJM, DRP, HMS, SAM, MLM, SM, TEP, TJS, DZ, CTB, and MJS). The U.S. Department of Agriculture supports Breeding Insight under agreement numbers [<u>8062-21000-</u> <u>043-004-A</u>, <u>8062-21000-052-002-A</u>, and <u>8062-21000-052-003-A</u>). Any opinions, findings, conclusions, or recommendations expressed in this publication are those of the Breeding Insight and do not necessarily reflect the views of the U.S. Department of Agriculture. In addition, any reference to specific brands or types of products or services does not constitute or imply an endorsement by the U.S. Department of Agriculture for those products or services.

### Clemson University

CC’s work on this project was made possible by the support of the American People provided to the Feed the Future Innovation Lab for Crop Improvement through the United States Agency for International Development (USAID) under Cooperative Agreement No. 7200AA-19LE-00005. BJE and TWR were supported by USDA NIFA SCRI No. 2022-51181-38449.

### BrAPI at Cornell University

The BrAPI Project and PS are funded by the USDA grant NIFA-DSFAS 2022-67021-37024.

### Boyce Thompson Institute

Funding for MFG and LAM was provided through a grant from the Gates Foundation (https://www.gatesfoundation.org/).

### Consultive Group on International Agricultural Research (CGIAR)

JSA was supported by the Alliance of Bioversity International and CIAT, part of CGIAR, a global research partnership for a food-secure future dedicated to transforming food, land, and water systems in a climate crisis. KA was supported by ICARDA, which is also part of CGIAR. JSA and KA would like to thank all funders who supported this research through their contributions to the CGIAR Trust Fund: https://www.cgiar.org/funders/.

## Notes

### Competing Interest Statement

The authors have declared no competing interest.

